# Finding informative neurons in the brain using Multi-Scale Relevance

**DOI:** 10.1101/316190

**Authors:** Ryan John Cubero, Matteo Marsili, Yasser Roudi

**Affiliations:** Kavli Institute for Systems Neuroscience and Centre for Neural Computation, Norwegian University of Science and Technology (NTNU), Olav Kyrres gate 9, 7030 Trondheim, Norway; The Abdus Salam International Center for Theoretical Physics, Strada Costiera 11, 34151 Trieste, Italy; Scuola Internazionale Superiore di Studi Avanzati, Via Bonomea 265, 34136 Trieste, Italy

## Abstract

We propose a metric – called Multi-Scale Relevance (MSR) – to score neurons for their prominence in encoding for the animal’s behaviour that is being observed in a multi-electrode array recording experiment. The MSR assumes that relevant neurons exhibit a wide variability in their dynamical state, in response to the external stimulus, across different time scales. It is a non-parametric, fully featureless indicator, in that it uses only the time stamps of the firing activity, without resorting to any *a priori* covariate or invoking any specific tuning curve for neural activity. We test the method on data from freely moving rodents, where we found that neurons having low MSR tend to have low mutual information and low firing sparsity across the correlates that are believed to be encoded by the region of the brain where the recordings were made. In addition, neurons with high MSR contain significant information on spatial navigation and allow to decode spatial position or head direction as efficiently as those neurons whose firing activity has high mutual information with the covariate to be decoded.

## Introduction

Next-generation techniques have allowed us to probe an increasing number of neurons in behaving animals [1]. Yet deciphering how specific functions are implemented in the neural code still remains a daunting task. In any case, the neurons that can be recorded are much less than those that are involved in the encoding of the animal’s behaviour; some of the recorded neurons may be attuned to different features of the stimuli or behaviour and some of them may display an activity that is not related to it. Much progress has been made in identifying those variations in the stimuli and the behaviours that correlate significantly with the firing pattern of individual neurons. Typical examples range from the discovery of simple and complex cells in the early visual cortices [2] to the more recent discovery of grid [3] and speed [4] cells. This approach has its limits: First, as observed experimentally e.g. by Sargolini *et. al* [3] and, more recently, using solid statistical analysis by Hardcastle *et al.* [5], the same neuron may respond to a combination of different behavioural covariates, such as position, head direction and speed in spatial navigation. Second, and most importantly, neurons may encode a particular behaviour in ways that are unknown to the experimenter and that are not related to covariates typically used or to *a priori* features. This has motivated a considerable amount of work in refining appropriate measures of covariates relevant for a particular behaviour. For example, the calculation for identifying grid cells, introduced in Ref. [3], has been refined in Ref. [6] and further in Ref. [7], in order to account for imperfect hexagonal symmetry of grid fields [8].

Here, we propose a novel non-parametric, model-free method for selecting relevant neurons that *does not require knowledge of external correlates*. This featureless selection is done by identifying neurons that have broad and non-trivial distribution of spike frequencies across a broad range of time scales. The proposed measure – called *Multi-Scale Relevance (MSR)* – allows the experimenter to rank the neurons according to their relevance to the behaviour probed in the experiment.

We illustrate the method by applying it to data on spatial navigation of freely roaming rodents in Refs. [9] and [10], that reports the neural activities of 65 neurons simultaneously recorded from the medial entorhinal cortex (mEC), and 746 neurons in the anterodorsal thalamic nucleus (ADn) and post-subiculum (PoS), respectively. In all cases, we find that neurons with low MSR also coincide with those that contain no information on covariates involved in navigation, but that the opposite is not true. Some neurons with high MSR also contain significant relevant information for spatial navigation, some relative to position, some to head direction but often on both space and head direction. This corroborates the recent conjecture that neurons in the mEC respond to a mix of features, rather than to a single one [5]. Furthermore, we show that the neurons in mEC with highest MSR have spike patterns that allow an upstream decoder “neuron” to discern the organism’s state in the environment. Indeed, the top most relevant neurons, according to MSR, decode spatial position (head direction) just as well as the top most spatially (head directionally) informative neurons.

MSR correlates to different degrees with different measures that have been introduced to characterise spatially specific neurons. For example, the MSR correlates strongly with spatial sparsity [11, 12] and weakly with grid score [3, 13]. Indeed, it has been proposed that grid score alone as defined in [3] is not efficient in identifying grid cells but its performance is improved when complemented with other measures [7]. This suggests that MSR can be used to pave a way towards identifying the external correlate which drive the activity of the neuron and finding those statistical features to which neurons are attuned to. Finally, a discussion on our findings and on possible future applications shall close the paper.

### Multi-Scale Relevance

We consider a population composed of *N* neurons in a freely behaving animal whose activities were simultaneously observed up to a time, *t_obs_*. The activity of neuron *i* is recorded and stamped by the spike times 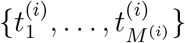 where 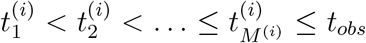 and *M* ^(*i*)^ is the total number of observed spikes of neuron *i*. We shall drop the superscript ^(*i*)^ henceforth when not needed, in order to simplify the notation. By discretizing the time into *T* bins of duration ∆*t*, a spike count code, *<u>s</u>* ={*k*_1_, *k*_2_, *…*, *k_T_*}, can be constructed where *k_s_* denotes the number of spikes recorded from neuron *i* in the *s*^th^ time bin *B_s_* = [(*s*–1)∆*t*, *s*∆*t*] (*s* = 1, 2, …, *T*).

Varying ∆*t* allows us to probe neural activity at different time scales. Yet, rather than using ∆*t* to measure time resolution, we adopt an information theoretic measure, given by

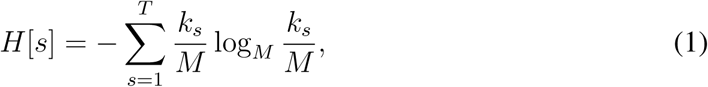

where log*_M_*(·) = log(·)/log *M* indicates logarithm base *M* (in units of *M* ats). Considering *k_s_/M* as the probability that neuron *i* fires in the bin *B_s_*, this has the form of a Shannon entropy [14]. This is the amount of information that one gains on the timing of a randomly chosen spike, by knowing the index *s* of the bin it belongs to [15]. We argue that *H*[*s*] provides an intrinsic measure of resolution, contrary to ∆*t* which refers to particular time scales that may vary across neurons. For example, there is a value 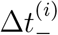 such that for all 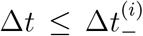 all time bins either contain a single spike or none, i.e. 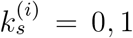 for all *s*. All these values of ∆*t* correspond to the same value of the intrinsic resolution *H*[*s*] = 1. Likewise, there may be a value 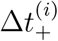 such that for all 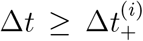 all spikes of neuron *i* fall in the same bin. All 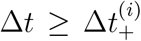 then correspond to the same value *H*[*s*] = 0 of the resolution, as defined here. In other words, *H*[*s*] captures resolution on a scale that is fixed by the available data.

Let us now move to a characterisation of the dynamic response of neuron *i*, at a given resolution *H*[*s*] (corresponding to a given ∆*t*). The only way in which the dynamic state of the neuron in bin *s* can be distinguished from that in bin *s′* is by its activity. If the number of spikes in the two bins is the same (*k_s_* = *k_s′_*) there is no way to distinguish the dynamic state of the neuron in the two bins, at that resolution [16]. Therefore, one way to quantify the richness of the dynamic response of a neuron is to count the number of different dynamic states it undergoes in the course of the experiment. A proxy of this is given by the variability of the spike frequency *k_s_*, that again can be measured in terms of an entropy

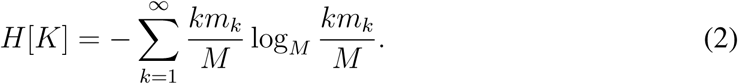

where *m_k_* indicates the number of time bins that contain *k* spikes [17], so that *km_k_*/*M* is the fraction of spikes that fall in bins with *k_s_* = *k*. Again, rather than considering *H*[*K*] as a Shannon entropy of an underlying distribution *p_k_* ≈ *km_k_*/*M* of spike frequencies, we take *H*[*K*] as an information theoretic measure of the information each spike contains on the dynamic state of the neuron at a given resolution [18]. Ref. [19] shows that *H*[*K*] measures the complexity of the variability in the sense that *H*[*K*] correlates with the number of parameters a model would require in order to describe properly the dataset, without overfitting. Hence, following Ref. [19], we shall call *H*[*s*] as *resolution* and *H*[*K*] as *relevance*.

In the current context, the reason for this choice can be understood as follows. In a freely behaving animal, different neurons can have activities that are more or less related to the behavioural states that are being probed in the experiment. Neurons that are *relevant* for encoding the animal’s behaviour are expected to display variation on a wide range of dynamical states, i.e. to have a large *H*[*K*]. On the contrary, neurons that are not involved in the animal’s behaviour are expected to visit relatively fewer dynamical states, i.e. to have a lower *H*[*K*].

Notice that for very small binning times 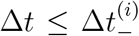 (when each time bins contains at most one spike, i.e. *m_k_*_=1_ = *M* and *m_k′_* = 0, ∀ *k′ >* 1) we find *H*[*K*] = 0 (and *H*[*s*] = 1). At the opposite extreme, when 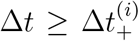 and *H*[*s*] = 0, we have all spikes in the same bin, i.e. *m_k_* = 0 for all *k* = 1, 2, …, *M* – 1 and *m_M_* = 1. Therefore again we find *H*[*K*] = 0. Hence, no information on the relevance of the neuron can be extracted at time scales smaller than 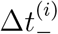 or larger than 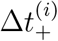. At intermediate scales 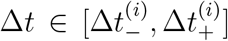, *H*[*K*] takes non-zero values [20], that we take as a measure of the relevance of neuron *i* for the freely-behaving animals being studied, at time scale ∆*t*.

Yet, the relevant time scale ∆*t* for a neuronal response to a stimulus may not be known *a priori* and/or the latter may evoke a dynamic response that spans multiple time scales. For this reason, we vary the binning time ∆*t* thereby inspecting multiple time scales with which we want to see the temporal code. As we vary ∆*t*, we can trace a curve in the *H*[*s*]- *H*[*K*] space for every neuron *i* in the sample. Neurons with broad distributions of spike frequencies across different time scales will trace higher curves in this space and in turn, will cover larger areas under this curve (see Fig. 1C). Henceforth, we shall call the area under this curve as the *multi-scale relevance* (MSR), ℛ_*t*_. The *relevant neurons*, those with high values of ℛ_*t*_, are expected to exhibit spiking behaviours that can be well-discriminated by upstream neurons over short and long time scales and thus, are expected to be *relevant* to the encoding of higher representations.

**Figure 1.**
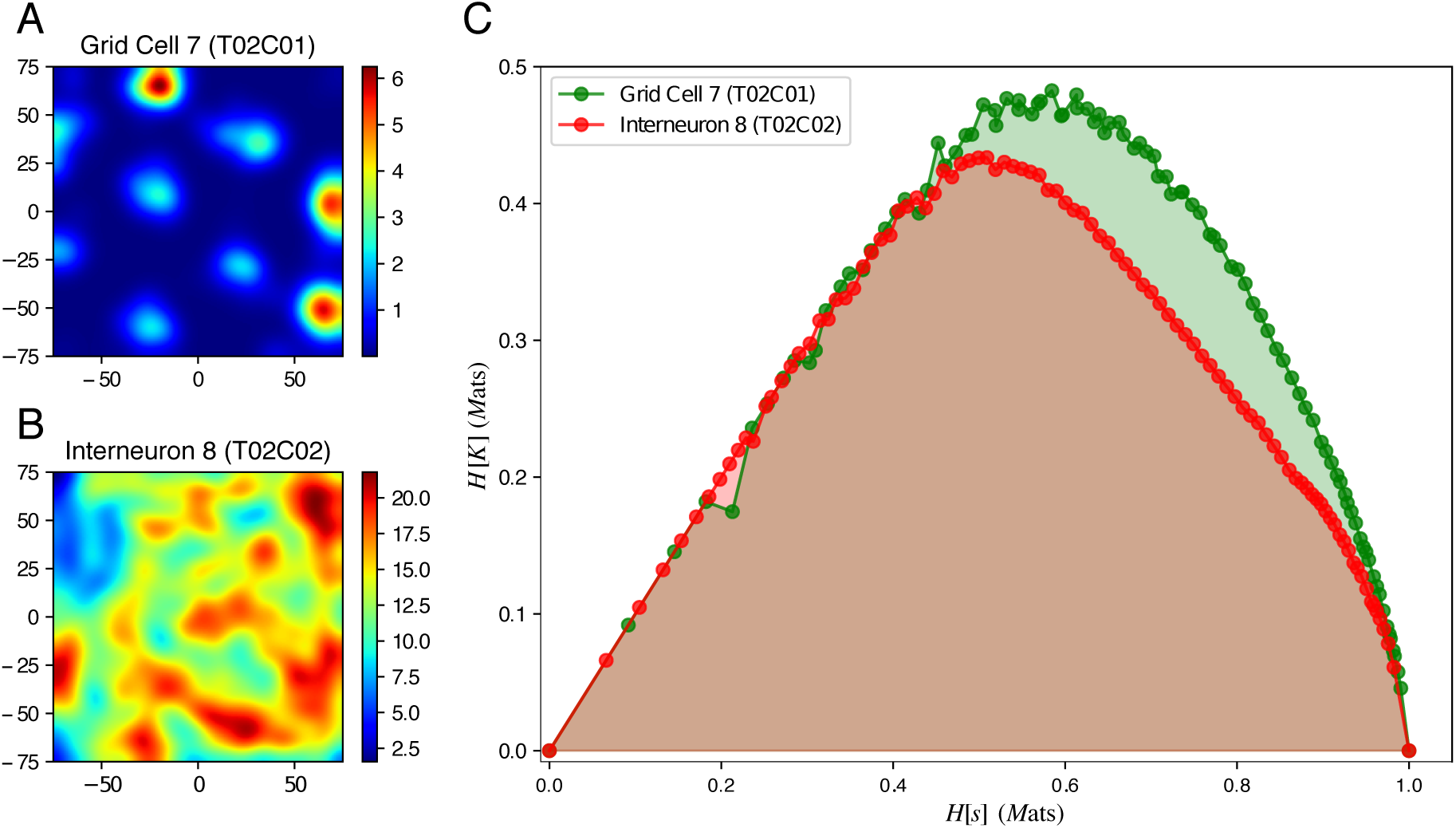
Proof of concept of the MSR as a relative information content measure. The smoothed firing rate maps of a grid cell (**A**) and an interneuron (**B**) in the mEC illustrates the spatial modulation of neural activity. Panel **C** shows the curves traced by the grid cell (green) and interneuron (red). Each point, (*H*[*s*], *H*[*K*]), in this curve corresponds to a fixed binning time, ∆*t*, with which we see the corresponding temporal neural spike codes.

MSR is designed to capture non-trivial structures in the spike time series. As such, it is expected to correlate with other measures characterising temporal structure, such as bursty-ness and memory [21]. Figure S1 shows indeed that, in synthetic data with given characteristics, MSR captures both the bursty-ness and memory of a time series. In addition, we find, in both synthetic and real data, a negative relation between MSR and spike frequency (i.e. *M*), which is partly associated with bursty-ness. Yet, the logarithm of the spike frequency (i.e., log *M*) cannot explain all of the variations in the MSR for real data which appears to be correlated with spatial and head directional information (see Figure S1).

As a proof of concept of the MSR for featureless neural selection, we considered two neurons recorded simultaneously from the medial entorhinal cortex (mEC) in Ref. [9] – a grid cell (T02C01) and an interneuron (T02C02) – both of which were measured from the same tetrode and thus, are in close proximity in the brain region. The mEC and its nearby brain regions are notable for neurons that exhibit spatially selective firing (e.g., *grid cells* and *border cells*) which provides the brain with a locational representation of the organism and provides the hippocampus with its main cortical inputs. Grid cells have spatially selective firing behaviours that form a hexagonal pattern which spans the environment where the rat freely explores as in Fig. 1A. Apart from spatial information, grid cells can also be attuned to the head direction especially in deeper layers of the mEC [3]. These cells altogether provide the organism with an internal map which it then uses for navigation. On the other hand, interneurons, as in Fig. 1B, are inhibitory neurons which are still important towards the formation of grid cell patterns [22, 23, 24] but have much less spatially specific firing patterns. Intuitively, as the mEC functions as a hub for memory and navigation, grid cells, which provide the brain with a representation of space, should be more relevant for an upstream “neuron” (possibly the place cells in the hippocampus) in encoding higher representations compared to interneurons. Indeed, the grid cell traces higher curves in the *Hs*] *− H*[*K*] space as in Fig. 1C and thus defines a larger area compared to the interneuron.

## Results

With the observations in Fig. 1, we sought to characterise the temporal firing behaviours of the 65 neurons which were simultaneously recorded from the mEC and its nearby regions of a male Long Evans rat as it freely explored a square arena of length 150 cm [9]. This heterogeneous neural ensemble, as characterised in Ref. [9], consisted of 23 grid cells, whose spatial firing fields can be clustered into 3 functional modules, 5 interneurons, 1 putative border cell and 36 unclassified neurons, some of which had highly spatially attuned firing behaviour and nearly hexagonal firing patterns [9, 25, 7]. This dataset was chosen among the multiple recording sessions performed in Ref. [9] as this contained the most grid cells to be simultaneously recorded.

These results were then corroborated by characterising the temporal firing behaviors of the 746 neurons which were recorded from multiple anterior thalamic nuclei areas, mainly the anterodorsal (AD) nucleus, and subicular areas, mainly the post-subiculum (PoS) of 6 different mice from across 31 recording sessions while the mouse explored a rectangular arena of dimensions 53 cm × 46 cm [10]. This data was chosen as these heterogeneous neural ensemble contained a number of *head direction cells* which are neurons that are highly attuned to head direction.

Before showing the results on these data sets, we note that the the MSR is a robust measure. To establish this, we compared the MSRs computed using only the first half of the data to that computed from the second half. As seen in Figure S2, we obtained very similar results, confirming that the MSR is a reliable measure that can be used to score neurons.

### MSR captures functionally relevant external correlates

As the mEC is crucial to spatial navigation, we expected that neurons with a high MSR score would contribute towards a representation of the animal’s spatial organization, in one way or another. Different measures relating the spatial position, x, with neural activity had been employed in the literature to characterise spatially specific neural discharges, like the Skaggs-McNaughton spatial information, *I*(*s*, x) defined in Eq. (7) and in Ref. [26], spatial sparsity measure, *sp*_x_ defined in Eq. (9) and in Ref. [11, 12] and grid score, *g*, defined in Eq. (10) and in Refs. [3, 13, 6, 7].

Apart from spatial location, head direction also plays a crucial role in spatial navigation. The mean vector length, *R* (Eq. (11) in Materials and methods) is commonly used as a measure of head directional selectivity of the activity of neurons. However, this measure assumes that there is only one preferred head direction in which a given neuron is tuned to. To this end, we calculated two measures – the head directional information, *I*(*s*, *θ*), and head directional sparsity, *sp_θ_* – inspired by the spatial information and spatial sparsity to quantify the information and selectivity of neural firing to head direction respectively. These measures ought to detect non-trivial and multimodal head directional tuning which may also be important in representing head direction in the brain [5].

In order to see whether spatially modulated firing behaviour can be captured solely from the spike times, as encoded in the MSR, Fig. 2 reports the Skaggs-McNaughton spatial information (A) and the analogous head directional information (D) as a function of the MSR for each neuron in the mEC data. Figs. 2B (C) and E (F) report the spatial firing rate maps and head direction tuning curves for the top ten (bottom ten) neurons by MSR score, respectively. This shows that neurons having a low MSR have very non-specific spatial and head directional discharges as indicated by their sparsity scores (Figs. 2C and F) whereas neurons having a high MSR have a broader range of spatial and head directional sparsity (Figs. 2B and E).

**Figure 2.**
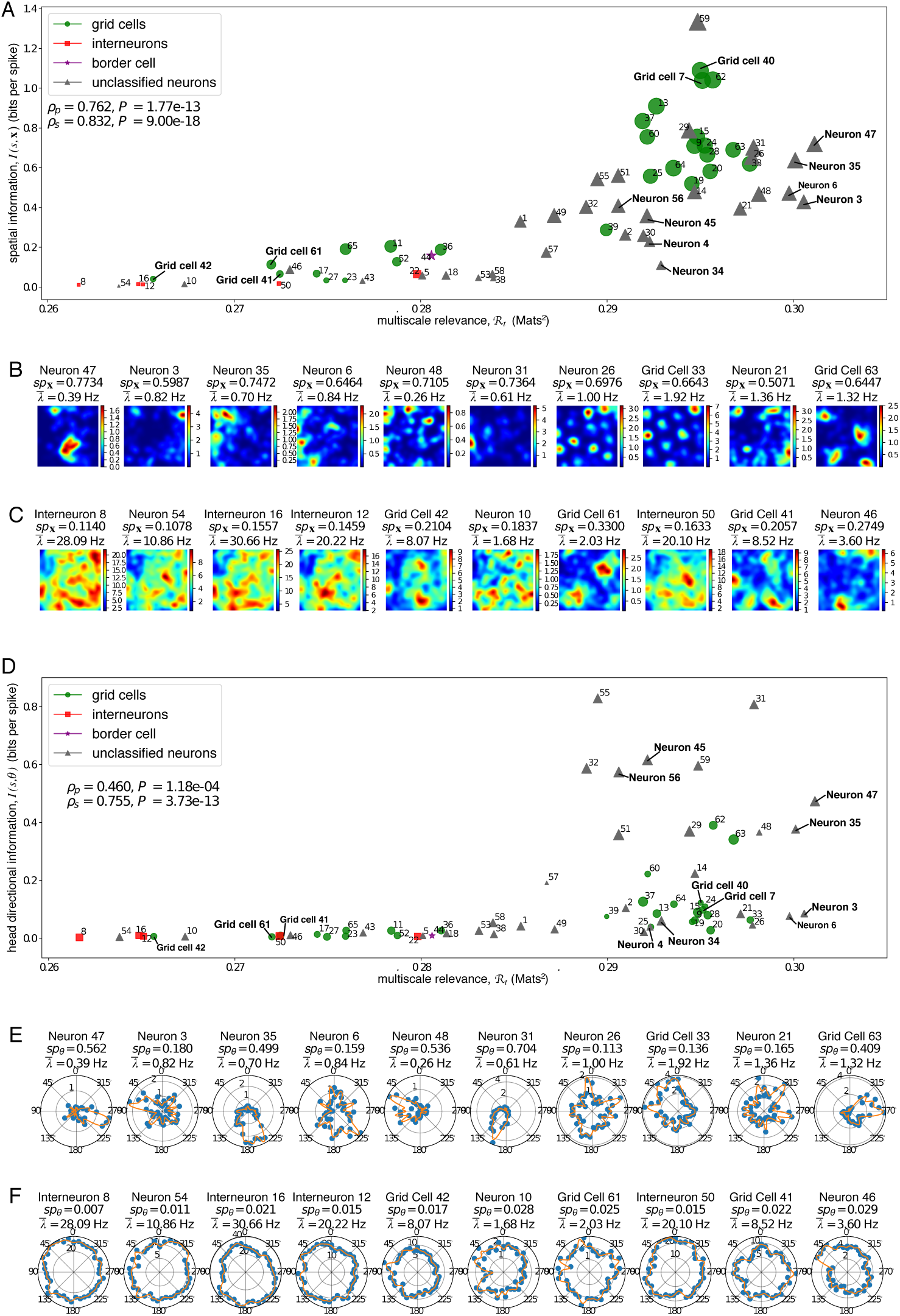
The multi-scale relevance identified neurons that are spatially and head directionally informative. A scatter plot of the MSR vs. the bias-corrected spatial (head direction) information is shown in **A** (**D**). The sizes of the scatter points reflect the spatial sparsity (head directional sparsity) of the neural activity while the shapes of the scatter points indicate the identity of the neuron according to Ref. [9]. The linearity and monotonicity of the multi-scale relevance and the information measures was assessed by the Pearson’s correlation, *ρ_p_*, and the Spearman’s correlation, *ρ_s_*, respectively. Information bias was measured by bootstrapping method, i.e., calculating the average of the spatial or head directional information of 1000 randomised spike trains. The spatial firing rate maps (head directional tuning curves) of the 10 most relevant neurons and the 10 most irrelevant neurons are shown together in panels **B** (**E**) and **C** (**F**) respectively with the calculated spatial sparsity, *sp*_x_, (head directional sparsity, *sp_θ_*) and mean firing rate, *λ̅*. Note that the mean firing rate, *λ̅*, was calculated using Eq. (8).

We found that (*i)* Neurons with high spatial information or high head direction information also had high MSR, but the converse was not true. While there are highly relevant neurons that responded exquisitely to space (grid cells 7 and 40) or head direction (neurons 45 and 56) alone, the majority (e.g. neurons 35 and 47) encoded significantly both spatial and head direction information. Secondly, we found that *ii*) Neurons with low MSR had both low spatial and low head direction information (Figs. 2C and F). Again, the converse was not true (e.g. neurons 4 and 34). Finally *iii*) we find that some neurons, for example, neurons 3 and 6, in spite of the fact that their rate maps (Figs. 2B and E) indicated some spatial and head directional sparsity, had relatively low spatial and head direction information but were both identified to be highly relevant neurons by MSR. This high MSR suggests that perhaps these neurons respond to different correlates involved in spatial navigation different from spatial location or head direction.

Many of the grid cells were spotted as highly relevant, but not all. For example, grid cells 41, 42 and 61, that had a significant grid score, had a low MSR (and low spatial information). This indicated that different measures correlate differently with MSR. Fig. 3 reports the distribution of the other four measures analysed in this study conditional to different levels of MSR. Fig. 3A shows that grid score maintains a large variation across all scales of the MSR, with a moderate increase in its average. A similar behaviour was observed in Fig. 3A for the mean vector length.

**Figure 3.**
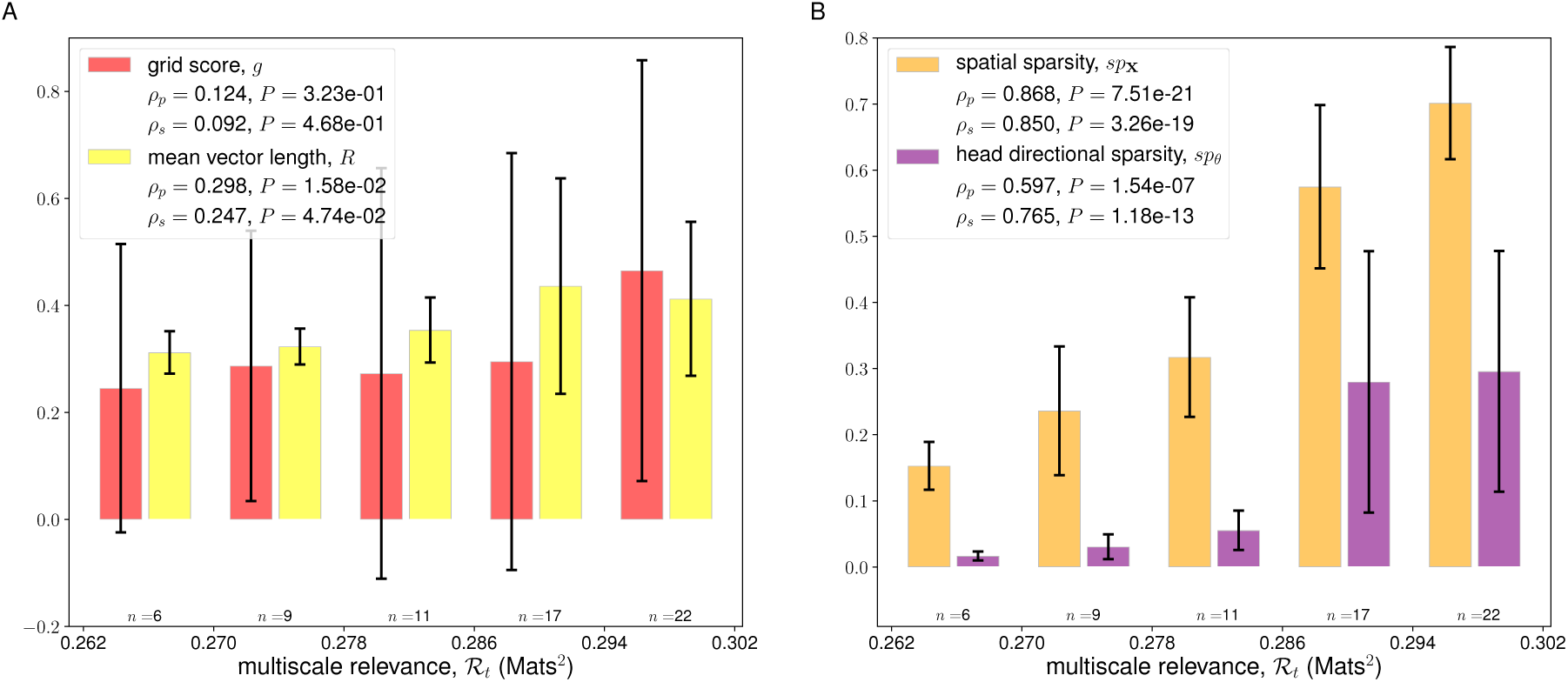
The MSR identified neurons with spatially and head directionally selective discharges. Bar plots depict the mean (height of the bar) along with the standard deviation (black error bars) of the grid score (red) and Rayleigh mean vector length (yellow) in panel **A**, and the spatial sparsity (orange) and head directional sparsity (purple) in panel **B** for each neuron in the mEC within the relevance range as indicated. The relevance range was determined by equally dividing the range of the calculated MSR into 5 equal parts. The number of neurons whose MSRs fall within a relevance range is indicated below each bar. The linearity and monotonicity between the multi-scale relevance and the different spatial and head directional quantities were quantified using the Pearson’s correlation, *ρ_p_*, and the Spearman’s correlation, *ρ_s_*, respectively.

Spatial sparsity and head directional sparsity, instead, showed a significant correlation with the MSR as seen in Fig. 3B. The observation that relevant neurons with high head directional sparsity may have low mean vector length was an indication that head directionally specific firing behaviour is not necessarily unimodal.

The ADn and PoS areas are known brain regions to contain head direction cells which robustly fires when the animal’s head is facing a specific direction [27, 28]. This network of head direction cells form part of the animal’s navigational system and is believed to be crucial to the formation of grid cells in the mEC [3, 29, 6]. Thus, we expected that head directionally attuned cells will be relevant to a freely behaving and navigating rodent. Indeed, as seen in Fig. 4, we observed that, in all of the 6 mice that were analysed, the head direction cells, i.e., neurons having high head direction sparsity and high mean vector lengths, had high MSRs. Focusing on a subset of neurons of Mouse 12 in Figs. 4A,B that were simultaneously recorded in a single session (Session 120806), we observed, as seen in Figs. 5A,B, that head directionally attuned neurons had high MSRs. However, the head direction alone may not explain the structure of the spike frequencies of these neurons. Hence, we also sought to find out whether some of these neurons are spatially tuned. As seen in Fig. 5E, we found that some of the relevant neurons were also modulated by the spatial location of the mouse.

**Figure 4.**
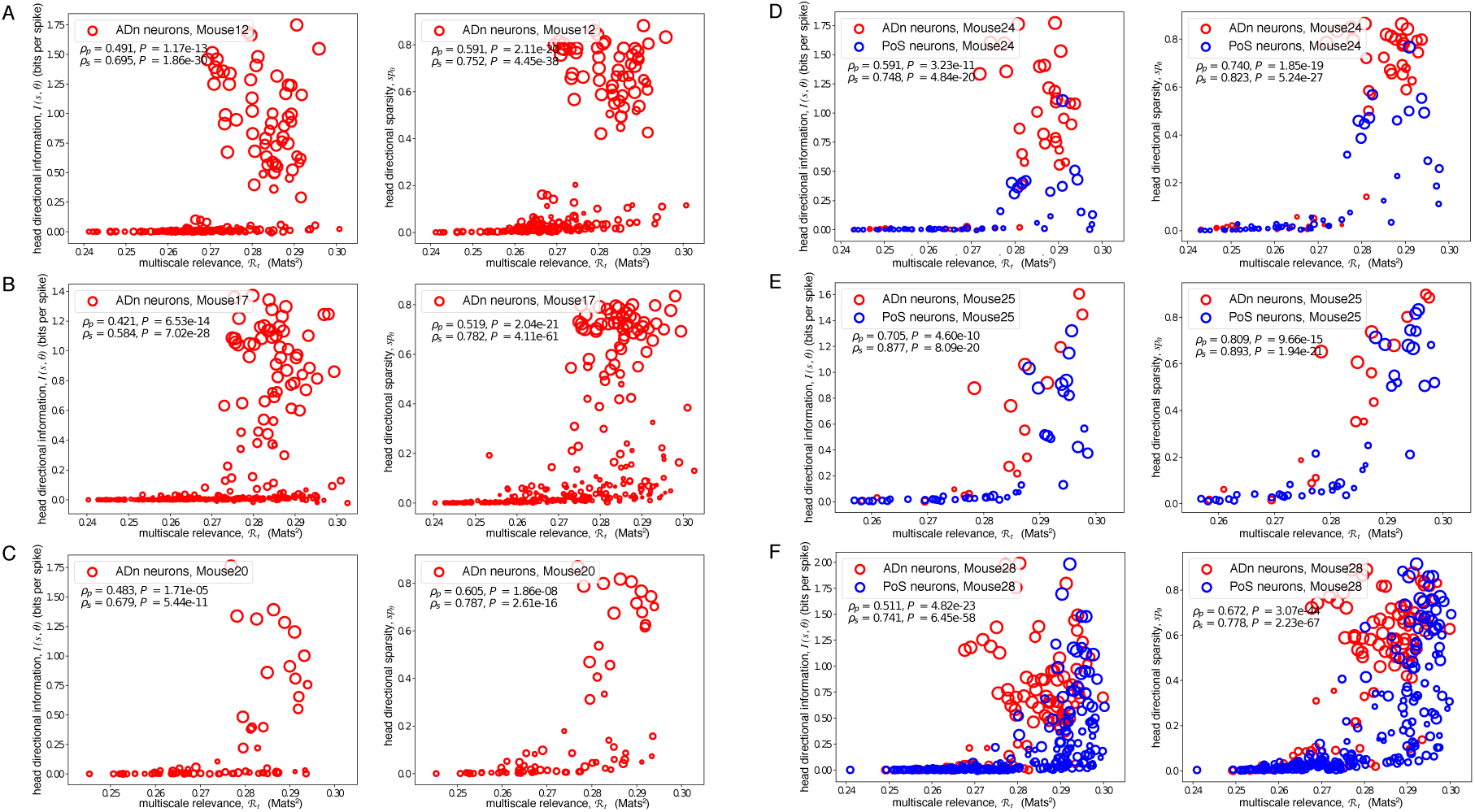
MSR of neurons from the anterodorsal thalamic nucleus (ADn) and post-subicular (PoS) regions of 6 freely-behaving mice pooled from multiple recording sessions. For each mice, the MSR of the recorded neurons which had more than 100 recorded spikes in a session were calculated. The corresponding the head directional information and sparsity (in bits per spike, see Materials and methods) were also calculated. ADn neurons are depicted in red circles while PoS neurons in blue circles. The size of each point reflect the mean vector lengths of the neurons wherein larger points indicate a unimodal distribution in the resulting head direction tuning curves.

**Figure 5.**
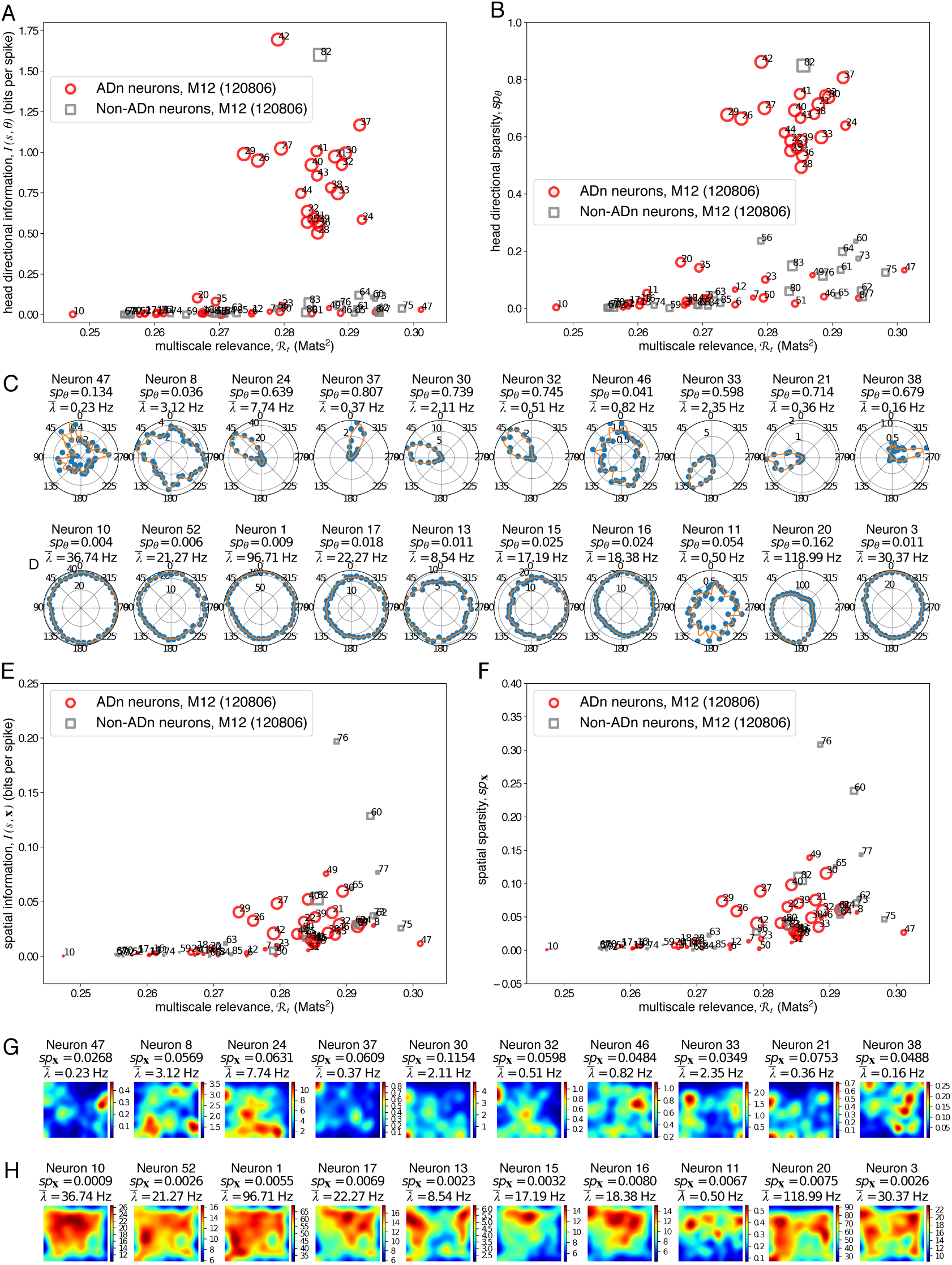
MSR of neurons from the anterodorsal thalamic nucleus (ADn) of Mouse 12 from a single recording session (Session 120806). A scatter plot of the multi-scale relevance vs. the bias-corrected head directional (spatial) information is shown in **A** (**E**). This plot is supplemented by a scatter plot between the multi-scale relevance and head directional (spatial) sparsity shown in **B** (**F**). The sizes of the scatter points reflect the mean vector length (head directional sparsity) of the neural activity where the larger scatter points correspond to putative head direction cells. The head directional tuning curves (spatial firing rate maps) of the 10 most relevant neurons and the 10 most irrelevant neurons are shown together in panels **C** (**G**) and **D** (**H**) respectively with the calculated head directional sparsity, *sp_θ_*, (spatial sparsity, *sp*_**x**_) and mean firing rate, *λ̅*).

To assess whether the spike frequencies, as characterised by the MSR, indeed contain information about external stimuli relevant to navigation, we resampled the spike count code of the neurons in the mEC such that only spatial information or only spatial and head directional information was incorporated. This resampling of the neural spiking was done by generating synthetic spikes assuming a non-homogeneous Poisson spiking with rates taken from the computed spatial firing rate maps and head directional tuning curves (See Materials and methods). These assumptions were able to recover the original rate maps as seen in Figs. 6C and D. Here, we focused our attention on *mEC Neuron 47* in the mEC data which had the highest MSR and also had both high spatial and high head directional information.

**Figure 6.**
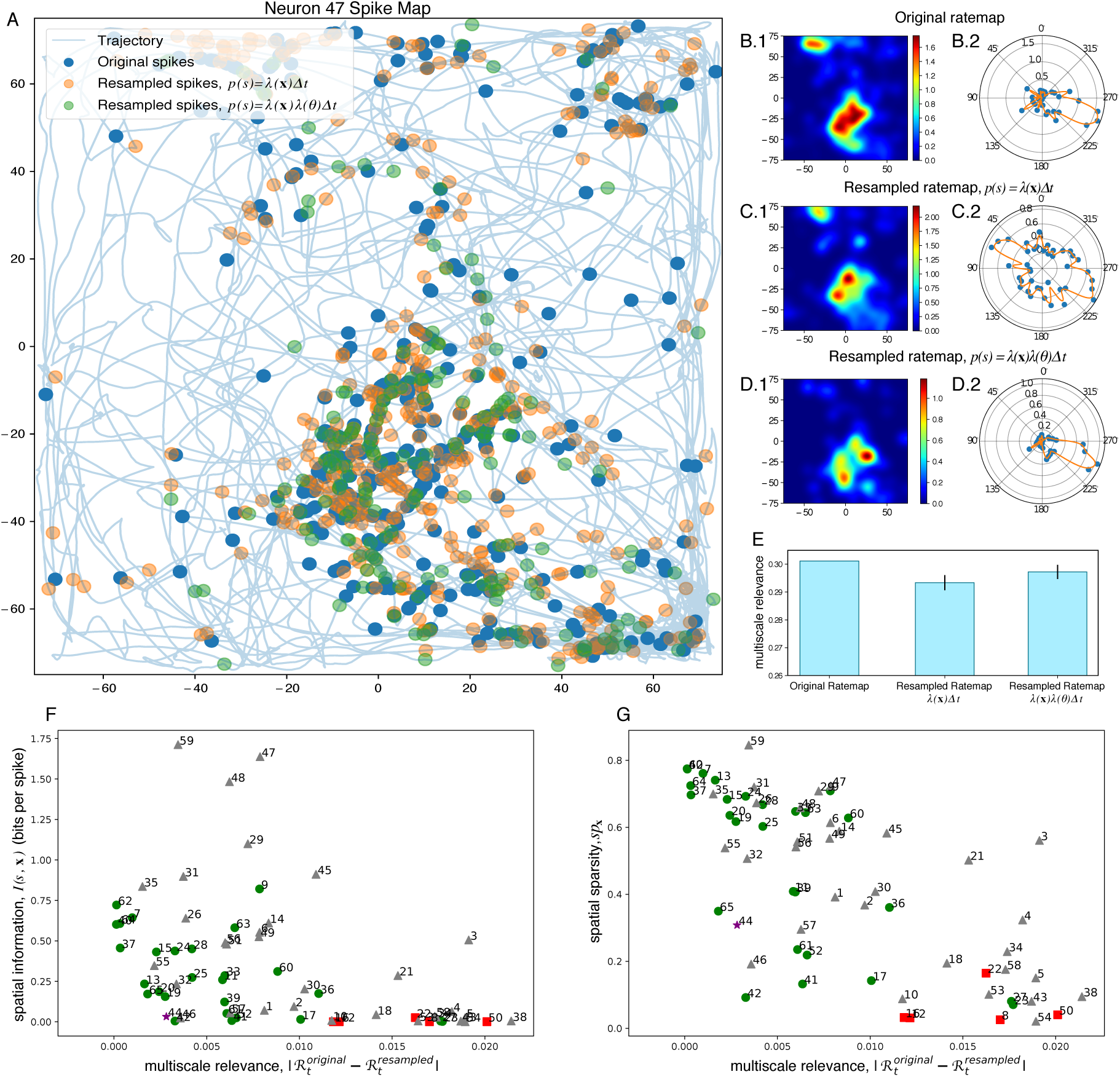
The MSR is a measure of information content of the neural activity. (**A**) Resampling the firing rate map using spatial position only (orange scatter points) or in combination with head direction (green scatter points) resulted to a firing activity that closely resembled the actual firing pattern (blue scatter points) of mEC Neuron 47. The blue lines indicate the real trajectory of the rat which was used when resampling the neural spiking. Compared to the original firing rate maps in **B**, the spatial (left panels) and head directional (right panels) firing rate maps were recovered by the resampling procedure in **C**-**D**. The result for a single realization of the resampling procedure is shown. (**E**) Bar plots show the multi-scale relevance calculated from the original spiking activity of the neuron and the resampled rate maps. The mean and standard deviation of 100 realizations of the resampling procedure is reported. Scatter plots between the difference of the multiscale relevance of the synthetic spikes and of the original spikes for each neuron and the spatial information (**F**), and the spatial sparsity (**G**) are also shown.

Indeed, by resampling only the spatial firing rate map as in Fig. 6E, we saw a decrease in the MSR despite having as much spatial information as the original spike count code. When head directional information was incorporated into the resampled spike frequencies, assuming the factorization of the firing probabilities due to position and head direction, more structure would be added onto the spiking activity of the resampled neuron. Hence, we expected to see an increase in the MSR as observed for *Neuron 47* which increased almost up to the MSR for the original spike frequencies. These findings support the idea that the temporal structure of the spike counts of the neuron, as measured by the MSR, come from its tuning profiles for both spatial position and head direction.

Furthermore, we also assessed which cells among the neurons in the mEC have MSRs that could be explained well by the spatial information and thus, were highly spatially attuned. We resampled the spatial firing rate maps of each of the cells in the mEC data (See Materials and methods). The absolute difference between the original and resampled MSR, 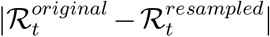, was then computed from the resampled spikes. When the variations in the spike frequencies could be explained by the spatial firing fields, we expected this difference to be close to zero. Indeed, as seen in Figs. 6F-G, we found that neurons having either high Skaggs-McNaughton spatial information or high spatial sparsity tended to have differential MSRs close to zero. Furthermore, we observed that most of the neurons having low differential MSRs were grid cells which have highly selective discharges with respect to the rat’s position.

Taken altogether, these analyses suggest that the MSR can be used to identify the interesting neurons in a heterogeneous ensemble. The proposed measure captures the non-trivial spike frequency distribution across multiple scales whose structure is highly influenced by external correlates that modulate the neural activity. Indeed, these analyses show that the MSR is able to capture information content of the neural spike code.

### Relevant neurons decode the positions as efficiently as spatially informative neurons

We found in the previous section that neurons with low MSR have low spatial or head directional information while higher MSR can indicate low or high values of spatial or head directional information. In this section, we show that despite this, high MSR can still be used to select neurons that decode position or head direction well. In other words, although high MSR can imply low spatial or head directional information, in terms of population decoding the set of high MSR neurons (selected based on only spike frequencies) performs equally well compared to the population of highly informative neurons (selected using the knowledge of the external covariate).

In order to understand whether MSR could identify neurons in mEC whose firing activity allows the animal to identify its position, we compared the decoding efficiency of the 20 neurons with the highest MSR with that of the 20 neurons with the highest Skaggs-McNaughton spatial information [26] (the two sets overlap on 14 neurons; see Fig. 2).

To this end, we employed a Bayesian approach to positional decoding wherein the estimated position at the *j*^th^ time bin, *x̂_j_*, is determined by the position, x_*j*_, which maximises an *a posteriori* distribution, *p*(x*_j_|*s_*j*_), conditioned on the spike pattern, s_*j*_, of a neural ensemble within the *j*^th^ time bin i.e.,

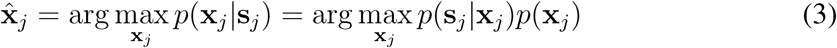

where the last term is due to Bayes rule, *p*(s*_j_|*x_*j*_) is the likelihood of a spike pattern, **s**_*j*_, given the position, **x**_*j*_, which depends on a given neuron model and *p*(x_*j*_) is the positional occupation probability which can be estimated directly from the data. Fig. 7A shows that a neural ensemble composed of relevant neurons decoded just as efficient as an ensemble composed of spatially informative neurons. It can also be observed that the relevant neurons decode the positions better than the ensemble composed solely of grid cells.

**Figure 7.**
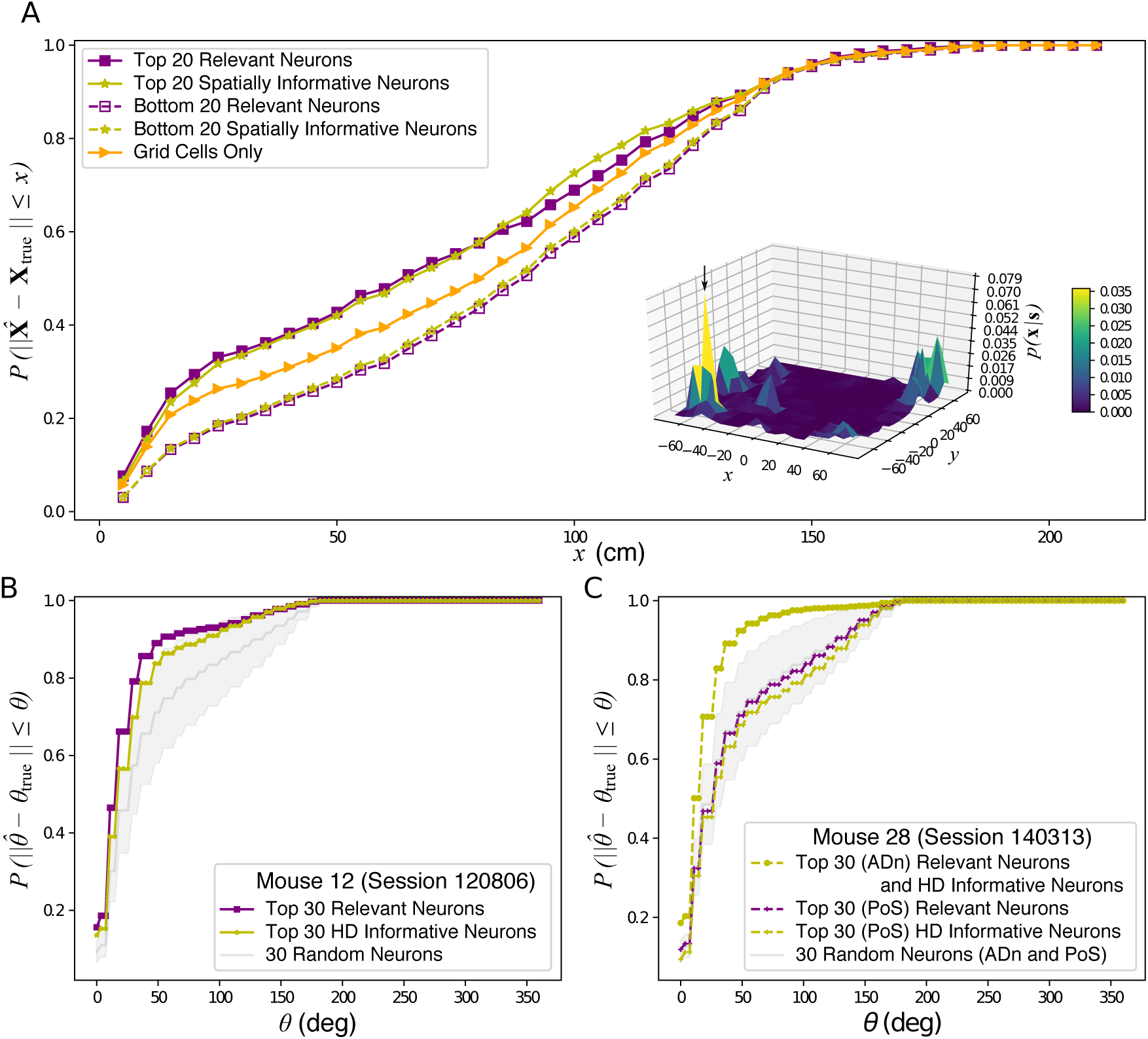
Positional decoding of relevant and informative neurons in the mEC and head directional decoding of the relevant and informative neurons in the ADn of Mouse 12 and the ADn and PoS of Mouse 28 under a single recording session. Panel **A** shows the cumulative distribution of the decoding error, ||**X**̂ – **X***_true_||*, for the relevant (solid violet) and spatially informative (solid yellow) neurons as well as for the irrelevant (dashed violet) and uninformative (dashed yellow) neurons. The low positional decoding efficiency at some time points can be traced to the posterior distribution, *p*(x**|**s), of the rat’s position given the neural responses which exhibited multiple peaks as shown in the inset surface plot. For this particular example, the true position was found close to the maximal point of the surface plot as indicated by the arrows although such was not always the case. Panel **B** depicts the cumulative distribution of the decoding errors of the 30 relevant (violet squares) and 30 head directionally informative (yellow stars) ADn neurons of Mouse 12 in Session 120806. The mean and standard errors of the cumulative distribution of decoding errors of 30 randomly selected ADn neuron (*n* = 1000 realizations) are shown in grey. On the other hand, panel **C** depicts the cumulative decoding error distribution of the 30 relevant (violet) and 30 head directionally informative (yellow) neurons in the ADn (crosses) and PoS (circles) of Mouse 28 in Session 140313. The mean and standard errors of the cumulative distribution of decoding errors of 30 randomly selected ADn or PoS neuron (*n* = 1000 realizations) are shown in grey. As the random selection included neurons from the ADn, which contain a pure head directional information and can decode the positions better than the neurons in the PoS, the decoding errors from the 30 randomly selected neurons were, on average, comparable to that of the relevant or head directionally informative PoS neurons. In all the decoding procedures, time points where all the neurons in the ensemble was silent were discarded in the decoding process.

Furthermore, we also took the 30 relevant and 30 head directionally informative ADn neurons from Mouse 12 (Session 120806) in Fig. 5 to decode for head direction. Mouse 12 was chosen as this animal had the most head direction cells recorded among the mice that only had recordings in the ADn [30]. In particular, we looked at the head direction decoding at longer time scales (in this case, ∆*t* = 100 ms), where we could model the neural activity using a Poisson distribution, *p*(n*_j_|θj*) similar to that in Eq. (15). Bayesian decoding adopts an equation

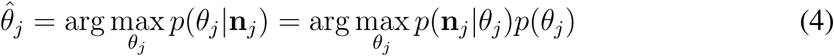

similar to Eq. (3) to estimate the decoded head direction, *θ̂_j_*, where *p*(*θ_j_*) is the head directional occupation as estimated from the data. We compared the decoding efficiency of the relevant neurons with the 30 ADn neurons with high head directional information which had 22 neurons that are relevant. We also compared the decoding efficiency of the 30 relevant or informative ADn neurons with 30 randomly selected ADn neurons (*n* = 1000 realizations). As seen in Fig. 7B, the 30 relevant neurons decoded just as well as the neural population composed of head directionally informative cells. Furthermore, the decoding efficiency of the relevant neurons were observed to be far better than the decoding efficiency of a random selection of neurons in the ensemble.

We also compared the decoding efficiency of the ADn and PoS neurons from Mouse 28 (Session 140313) which had the most head direction cells recorded among the mice that had recordings in both ADn and PoS [30]. As seen in Fig. 7C, neurons in the ADn decoded the head direction more efficiently than the neurons in the PoS. These results are consistent with the notion that the ADn contains pure head directional modulation which allow for head direction cells in the ADn to better predict the head direction compared to the head direction cells in the PoS which contain true spatial information [10, 31]. For the neurons in Mouse 28 (Session 140313), it had to be noted that the 30 relevant ADn neurons also happened to be the 30 head directionally informative ADn neurons. On the other hand, among the 30 relevant PoS neurons, 23 of which were head directionally informative. We observed that the relevant PoS neurons decode just as efficient as the informative neurons consistent with the findings for Mouse 12 (Session 120806).

Taken altogether, despite being blind to the rat’s position and of the mouse’s head direction, the MSR is able to capture neurons that can decode the position and head direction just as well as the spatially informative neurons and as the head directionally informative neurons.

## Discussion

In the present work, we introduced a novel, parameter-free and fully featureless method – which we called multi-scale relevance (MSR) – to select relevant neurons within a heterogeneous population of neurons which are supposed to respond to some external stimuli or to encode the behaviour of an animal that is being observed in an experiment. In the task of learning from the temporal code, we have shown that the information provided by the MSR can be compared to different correlates and allows one to disentangle the responses across stimuli or behaviours, some of which may also be unknown to the experimenter *a priori*.

In this paper, we assumed that the information carried by the activity of a given neuron is encapsulated in the long-ranged statistical patterns of the spike activity. In order to quantify this information, we used the ideas in Refs. [19] and [32] to hypothesise that neurons having such non-trivial temporal structures, as manifested by broad distributions of the neural firing behaviour, are important to the representations that the brain region encodes. At a given resolution, as defined in Eq. (1), we estimate the complexity of the temporal code by the relevance defined in Eq. (2). The latter captures the broadness of the spike frequency distribution at that resolution. Since naturalistic and dynamic stimuli and behaviours often operate on multiple time scales, the MSR integrates over different resolution scales, thus allowing us to spot neurons exhibiting persistent non-trivial spike codes across a broad range of time scales.

Here, we have shown that the neurons showing persistently broad spike frequency distributions across a wide range of time scales usually carry information about the external correlates related to the behaviour of the observed animal. By analysing the neurons in the mEC and nearby brain regions and the neurons in the ADn and PoS – brain structures that are pertinent to spatial navigation – we showed that the relevant neurons in these regions have firing behaviours that are selective to spatial location and head direction. Here, we found that in many cases, the neurons that display broad spike distributions tend to have conjugated representations in that they exhibit high mutual information with multiple behavioural features. These findings are consistent with those observed experimentally by Sargolini *et. al* [3] and statistically by Hardcastle *et al.* [5].

Broad distributions of spike frequencies, characterised by a high MSR, exhibit a stochastic variablility that requires richer parametric models, as shown in Ref. [19]. In a decoding perspective, these non-trivial distributions afford a higher degree of distinguishability of neural responses to a given stimuli or behaviour. Indeed, by decoding for either spatial position or for head direction using statistical approaches, we found that the responses of high MSR neurons allow upstream processing units to efficiently decode the external correlates just as well as the neurons whose resulting tuning maps contain information about those external correlates.

Finally, we observed that the population of relevant neurons, as identified by the MSR, is not homogeneous, e.g., the relevant neurons in the mEC data are not composed solely by grid cells and the relevant neurons in the ADn and PoS are not necessarily composed solely of head directional cells. Noteworthy, the decoding efficiency of the relevant neurons was observed to be better compared to the ensemble comprising solely the grid cells. When taken altogether, these observations support the idea that population heterogeneity may play a role towards efficient encoding of stimuli [33, 34].

The insistence on broad distributions, on which the MSR relies on, tails with the fact that biological systems such as the one under study, hardly ever generates well sampled datasets of their complex behaviour. The dynamical range which the experimenter can probe is limited by the size of the dataset and its often far from saturating biological dynamical ranges. The MSR takes advantage of this feature and identifies those variables that exhibit a richer variability. This intuition, discussed theoretically in Refs. [19] and [32], has also been used to identify biologically and evolutionary relevant amino acid sites in protein sequences [35]. Indeed, in spite of all advances in sequencing techniques, the genomes from which we can learn are only those left to us by evolution. Hence, Ref. [35] shows that subsequences that exhibit a wider response in frequency – as measured by Eq. (2) – to evolutionary dynamics contain a wealth of biologically relevant information. This same strategy can be used to identify relevant and hidden (latent) variables in statistical learning (see e.g. [36, 37]).

In principle, multi-scale relevance can be extended to measure the information carried by the spike time series with respect to known external correlates or features, such as spatial location and head direction. This requires discretising the feature space (e.g. space) in bins and computing the number of time with which a neuron fires in each bin. From the distribution of these counts, one can derive a measure of resolution and relevance, as in Eqs. (1,2), and draw a curve as in Fig. 1 upon changing the bin size. Like the MSR discussed here, this multi-scale relevance would be designed to spot neurons having spike frequencies with non-trivial distributions when projected onto the feature space. Indeed, Refs. [7] and [41] has shown that grid cells have field-to-field variability which is robust and is not an artifact of having finite data nor of the non-uniform spatial sampling and which has multiple implications including the capacity of grid cells to contain contextual information contrary to the findings in Refs. [38, 39, 40] as well as the remapping of place cells without the need for changing grid cell phases [41]. Although the application of MSR adapted to space to characterise spatial inhomogeneity of firing behaviour is an interesting avenue of further research, here we limited our analysis to temporal binning precisely because it is defined in terms of the sole spike activity – the only information available to upstream neurons – to decode a representation of the feature space.

The fact that the MSR captures these functional information from the temporal code is a remarkable feat of this measure. This method can then be used as a pre-processing tool to impose a less stringent criteria compared to those widely used in many studies (e.g., mean vector length, spatial sparsity and grid scores) thereby directing further investigation to interesting neurons. The MSR is expected to be particularly useful in detecting relevant neurons in high-throughput studies where the activity of thousands of neurons are measured and where the function of these neural ensemble are not known *a priori*. Whether this measure can also be used to identify functionally relevant neuronal units recorded through calcium imaging or through fMRI is also an exciting direction for future studies.

## Materials and methods

### Data Collection

The data used in this study are recordings from rodents with multisite tetrode implants. These neurons are of particular interest because they are involved in spatial navigation.

#### Data from medial entorhinal cortex (mEC)

The spike times of 65 neurons recorded across the mEC area of a male Long Evans rat (Rat 14147) were taken from Ref. [9]. The rat was allowed to freely explore a box of dimension 150 × 150 cm^2^ for a duration of around 20 mins. The positions were tracked using a platform attached to the head with red and green diodes fixed at both ends. Additional details about the data acquisition can be found in Ref. [9].

#### Data from the anterodorsal thalamic nucleus (ADn) and post-subiculum (PoS)

The spike times of 746 neurons recorded from multiple areas in the ADn and PoS across multiple sessions in six free moving mice (Mouse 12, Mouse 17, Mouse 20, Mouse 24, Mouse 25 and Mouse 28) while they freely foraged for food across an open environment with dimensions 53 × 46 cm^2^ and in their home cages during sleep were taken from Ref. [30]. Mouse 12, Mouse 17 and Mouse 20 only had recordings in the ADn while Mouse 24, Mouse 25 and Mouse 28 had simultaneous recordings from ADn and PoS. The positions were tracked using a platform attached to the heads of the mice with red and blue diodes fixed at both ends. Only the recorded spike times during awake sessions and the neural units with at least 100 observed spikes were considered in this study. Additional information regarding the data acquisition can be found in Refs. [10] and [30].

### Position and speed filtering

The position time series for the mEC data were smoothed to reduce jitter using a low-pass Hann window FIR filter with cutoff frequency of 2.0 Hz and kernel support of 13 taps (approximately 0.5 s) and were then renormalised to fill missing bins within the kernel duration as done in Ref. [7]. The rat’s position was taken to be the average of the recorded and filtered positions of the two tracked diodes. The head direction was calculated as the angle of the perpendicular bisector of the line connecting the two diodes using the filtered positions. The speed at each time point was computed by dividing the trajectory length with the elapsed time within a 13-time point window. When calculating for spatial firing rate maps and spatial information (see below), only time points where the rat was running faster than 5 cm/s were considered. No speed filters were imposed when calculating for head directional tuning curves and head directional information. On the other hand, no position smoothing nor speed filtering were performed when calculating for the spatial firing rate maps and spatial information for the ADn and PoS data.

### Rate maps

The spike location, 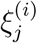, of neuron *i* at a spike time 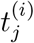 was calculated by linearly interpolating the filtered position time series at the spike time. As in Ref. [7], the spatial firing rate map at position x = (*x*, *y*) was calculated as the ratio of the kernel density estimates of the spatial spike frequency and the spatial occupancy, both binned using 3 cm square bins, as

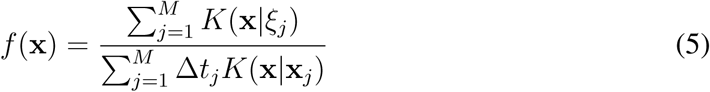

Where a triweight kernel

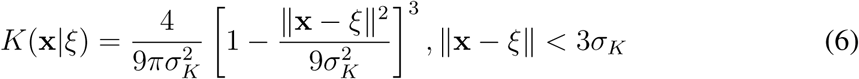

with bandwidth *σ_K_* = 4.2 cm was used. In place of a triweight kernel, a Gaussian smoothing kernel with *σ_G_* = 4.0 truncated at 4*σ_G_* was also used to estimate the rate maps which gave qualitatively similar results. For better visualization, a Gaussian smoothing kernel with *σ_G_* = 8.0 was used to filter the spatial firing rate map.

On the other hand, for head direction tuning curves, the angles were binned using 9*°* bins. The tuning curve was then calculated as the ratio of the head direction spike frequency and the head direction occupancy without any smoothing kernels as the head direction bins are sampled well-enough. For better visualization, a Gaussian kernel with smoothing window of 20*°* was used to filter the tuning curves.

### Information, Sparsity and other Scores

Given a feature, *ϕ* (e.g., spatial position, x, head direction, *θ* or speed, *υ*), the information between the neural spiking s and the feature can be calculated á la Skaggs-McNaughton [26]. In particular, under the assumption of a non-homogeneous Poisson process with feature dependent rates, *λ*(*ϕ*), under small time intervals ∆*t*, the amount of information, in bits per second, that can be decoded from the rate maps is given by

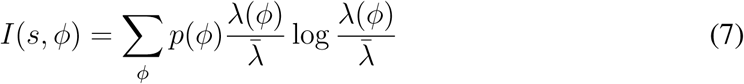

where *λ*(*ϕ*) is the firing rate at *ϕ*, *p*(*ϕ*) is the probability of occupying *ϕ* and

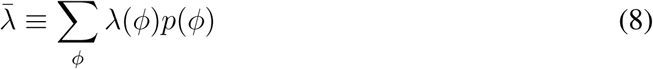

is the average firing rate. To account for the bias due to finite samples, the information of a randomised spike frequency was calculated using a bootstrapping procedure. To this end, the spikes were randomly shuffled 1000 times and the information for each shuffled spikes was calculated. The average randomised information was then subtracted from the non-randomised information.

Apart from the information, one of the measures that are used to quantify selectivity of neural firing to a given feature is the firing sparsity [12] which can be calculated using

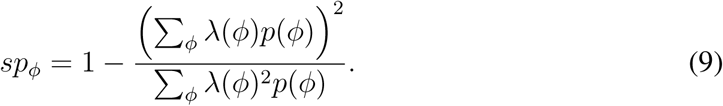

Apart from the measures of information and sparsity, we also calculated the grid scores, *g*, for the neurons in the mEC data. The grid score is designed to quantify the hexagonality of the spatial firing rate maps through the spatial autocorrelation maps (or autocorrelograms) and was first used in Ref. [3] to identify putative grid cells. In brief, the grid score is computed from the spatial autocorrelogram where each element *ρ_ij_* is the Pearson’s correlation of overlapping regions between the spatial firing rate map shifted *i* bins in the horizontal axis and *j* bins in the vertical axis and the unshifted rate map. The angular Pearson autocorrelation, acorr(*u*), of the spatial autocorrelogram was then calculated using spatial bins within a radius *u* from the center at lags (or rotations) of 30*°*, 60*°*, 90*°*, 120*°* and 150*°*, as well as the ±3*°* and ±6*°* offsets from these angles to account for sheared grid fields [8]. As done in Ref. [7], the grid score, *g*(*u*), for a fixed radius of *u*, is computed as

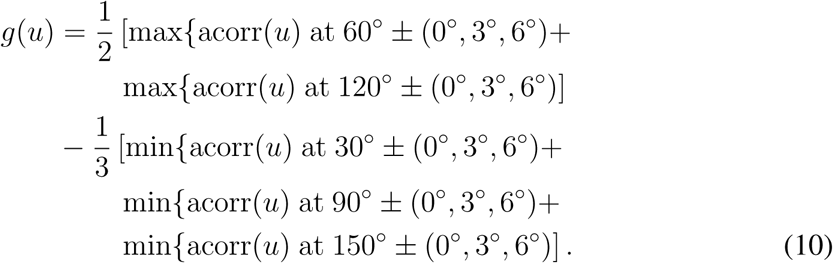

The final grid score, *g*, is then taken as the maximal grid score, *g*(*u*), within the interval *u* ∈ [12 cm, 75 cm] in intervals of 3 cm.

Another quantity that was calculated in this paper is the Rayleigh mean vector length, *R*. Given the angles {*θ*_1_, …, *θ_M_*} where a neuronal spike was recorded, the mean vector length can be calculated as

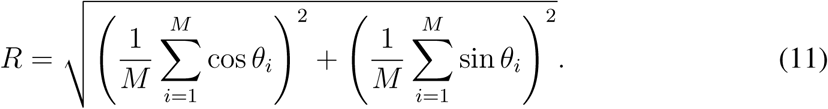

Note that for head direction cells where the neuron fires at a specific head direction, the angles will be mostly concentrated along the preferred head direction, *θ_c_*, and hence, *R ≈* 1 whereas for neurons with no preferred direction, *R ≈* 0.

### Resampling the firing rate map

The calculated rate maps and the real animal trajectory were used to resample the neural activity assuming non-homogeneous Poisson spiking statistics with rates taken from the rate maps. To this end, the real trajectory of the rat was divided into ∆*t* = 1 ms bins. The position and head direction were linearly interpolated from the filtered positions described above. The target firing rate, *f_j_* in bin *j* was then calculated by evaluating the tuning profile at the interpolated position or head direction. Whenever the target firing rate was modulated by both the position and head direction, we assumed that the contribution due to each feature was multiplicative and thus, *f_j_* is calculated as the product of the tuning profiles at the interpolated position and the interpolated head direction. A Bernoulli trial was then performed in each bin with a success probability given by *f_j_*∆*t*.

### Statistical decoding

For positional decoding, we divided the space in a grid of 20×20 cells of 7.5 cm × 7.5 cm spatial resolution, which was comparable to the rat’s body length. Time was also discretized into 20 ms bins which ensured that for most of the time (i.e. in 92% of the cases), the rat was located within a single spatial cell. Under these time scales, the responses of a neuron can be regarded as being drawn from a binomial distribution, i.e., either the neuron *i* is active 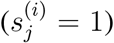 or not 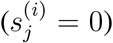 between (*j −* 1)∆*t* and *j*∆*t*. The likelihood of the neural responses, 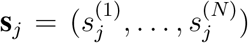 of *N* independent neurons at a given time conditioned on the position, x*_j_* is then given by

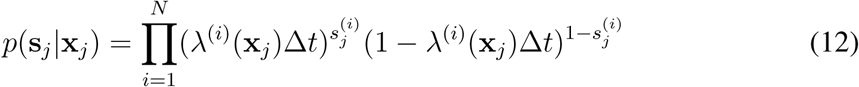

where *λ*^(*i*)^(x_*j*_) is the firing rate of neuron *i* at x_*j*_. Given the prior distribution on the position, *p*(x_*j*_), which is estimated from the data, the posterior distribution of the position, x_*j*_, given the neural responses, s*_j_* at time *t* is given by

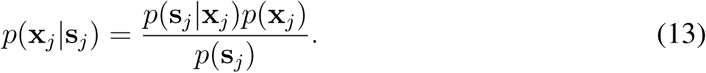

The decoded position, as in the Bayesian 1-step decoding in Ref. [42], was calculated as

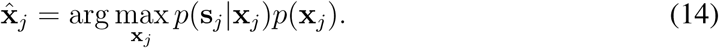

For head directional decoding, on the other hand, we divided the angles, *θ ∈* [0, 2*π*) in 9*°* bins. For this case, time was instead discretised into 100 ms bins. Under these time scales, the neurons could not be regarded simply as either active or not. Hence, it was natural to switch towards the analysis of population vectors, n_*j*_, a vector which represents the number of spikes, 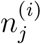, recorded from each neuron within the *j*^th^ time bin, to decode for the head direction. In this case, the number of spikes, 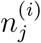, that neuron *i* discharges between (*j–*1)∆*t* and *j*∆*t* can be modeled as a non-homogeneous Poisson distribution

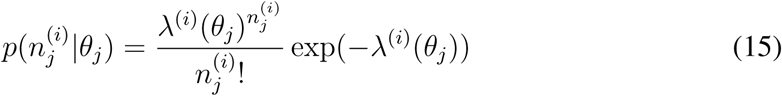

and thus, under the independent neuron assumption, 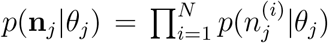. The decoded head direction can then be calculated as

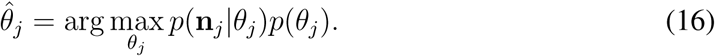

where *p*(*θ_j_*) is the head directional prior distribution which is estimated from the data. Note that in all of the decoding procedures, we only decoded for time points with which at least one neuron was active.

### Source codes

All the calculations in this manuscript were done using personalised scripts written in Python 3. The source codes for calculating multi-scale relevance (which is also compatible with Python 2) and for reproducing the figures in the main text are accessible online at https://github.com/rcubero/MSR.

## Supporting information

**Figure S1.**
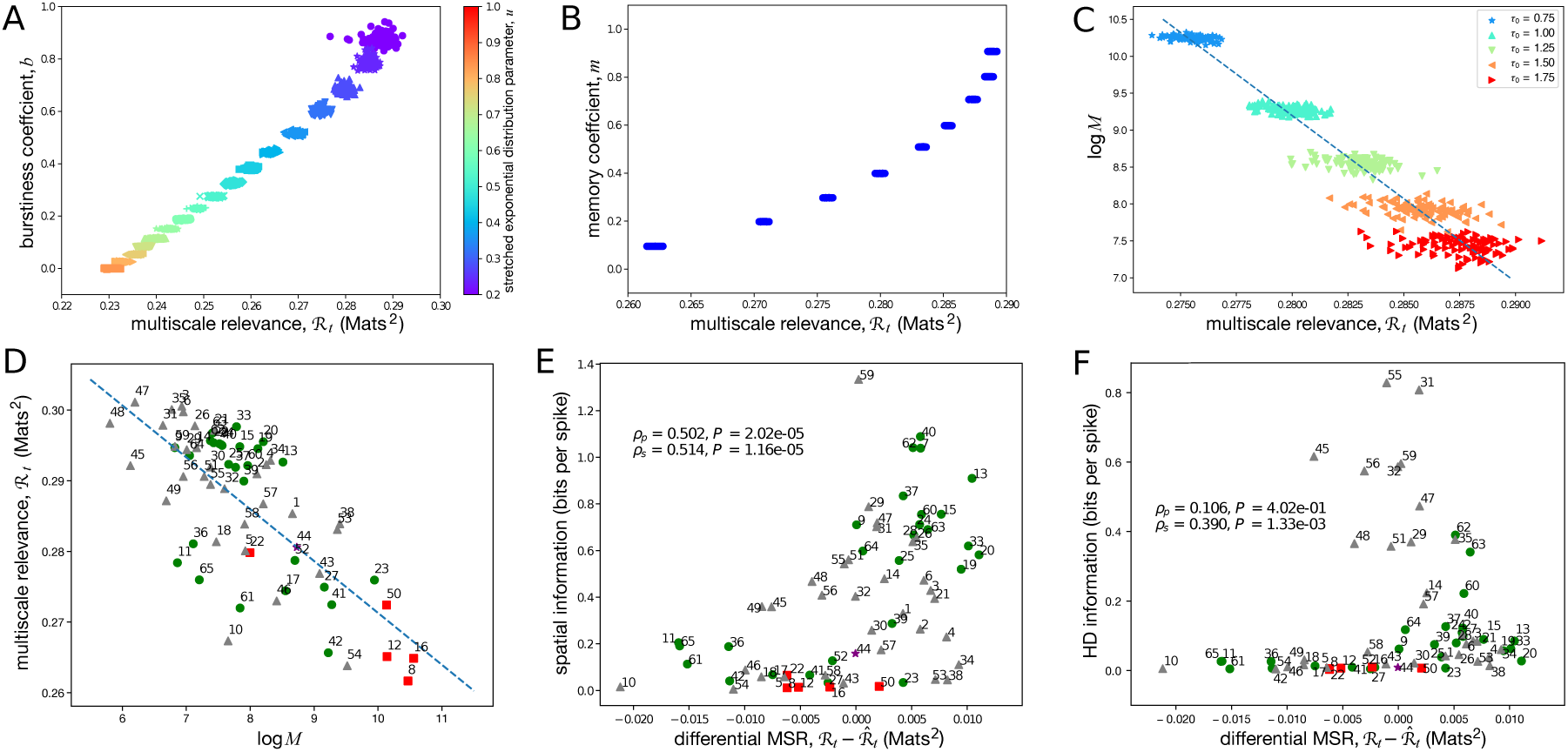
Relationship between the MSR and the bursty-ness and memory coefficients for synthetic data and neural data. Interevent times were drawn from a stretched exponential distribution to simulate random events up to 100,000 time units where shortterm memory effects were introduced through a shuffling procedure and the number of random events, *M*, were varied by modifying the characteristic time constant, *τ*_0_ (See Text S1 for details). Scatter plots show how the multi-scale relevance (MSR) scales with the bursty-ness coefficient, *b* (panel **A**), the memory coefficient, *m* (panel **B**), and log *M* (panel **C**). In panel **B**, random events were drawn from a stretched exponential distribution with *u* = 1.0 while in panel **C**, the parameter *u* was set to 0.3. The results for 100 realizations of such random events are shown. For the neurons in the mEC dataset, the MSR was linearly regressed with log *M* (panel **D**). The residuals, defined as the deviation of the MSR from the black dashed line, were then correlated against spatial (panel **E**) and head directional (panel **F**) information.

**Figure S2.**
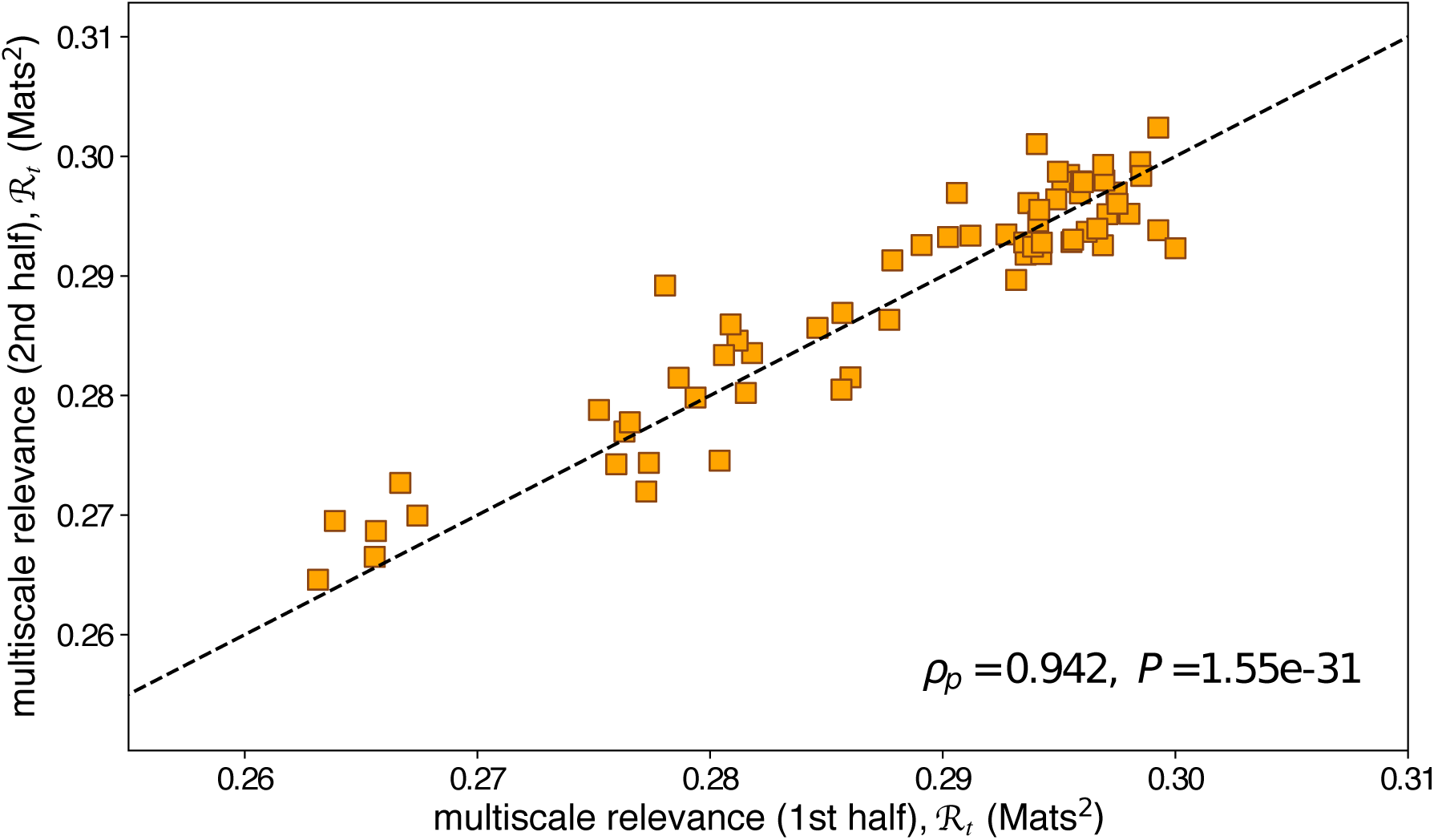
The multi-scale relevance is robust and it remains consistent even when using partial data. For each neuron, the multi-scale relevance was calculated using only the first half and only the second half of the data. The scatter plot reports the two results. The linearity of the relationship between the two sets of partial data is quantified by the Pearson correlation *ρ_p_* along with its *P*-value. The black dashed line indicates the linear fit.

**Text S1** Probes the relation between MSR and other measures of temporal structure using synthetic neural spiking.

## Text S1 Relation between MSR and other measures of temporal structure

Characterising the neural spiking can be done by studying the distribution of the time intervals between two succeeding spikes, known in literature as the interspike interval (ISI) distribution which allows us to see whether a neuron fires in bursts [1, 2]. Note that given the time stamps of neural activity {*t*_1_, …, *t_M_*}, the interspike interval is given by {*τ*_1_, …, *τ_M_*_−1_} where *τ_i_* = *t_i_*_+1_–*t_i_*. Because the MSR is built to separate relevant neurons from the irrelevant ones through their temporal structures in the neural spiking, we wanted to assess how the proposed measure scales with the characteristics that give structure to temporal events. In the context of the temporal activity of a neuron, a feature of the relevance measure, *H*[*K*] is that highly regular, equally-spaced interspike intervals are attributed with a low measure. On the other hand, interspike intervals that follow broad, non-trivial distributions are attributed with a high relevance measure. Hence, we expected that the relevance measure, and therefore the multi-scale relevance, captures non-trivial bursty patterns of neurons.

To study how MSR behaves with respect to the characteristics of interspike intervals, we considered a stretched exponential distribution

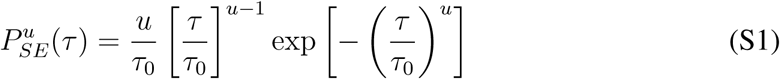

with which the parameter *u* allows us to define the broadness of underlying distribution and *τ*_0_ is the characteristic time constant of the random event. For Poisson processes, the interspike intervals follow an exponential distribution corresponding to *u* = 1 in Eq. (S1). For *u <* 1, the interspike interval distribution becomes broad and tends to a power law distribution with an exponent of –1 in the limit when *u→*0. On the other hand, for *u >* 1, the distribution becomes narrower and tends to a Dirac delta function in the limit when *u→∞*.

Upon fixing the parameters *u* and *τ*_0_ which fixes the stretched exponential distribution in Eq. (S1), random interspike intervals *τ_i_* could then be sampled independently from Eq. (S1) so as to generate a time series of 100,000 time units. The MSRs of each time series could then be calculated using the methods described in the main text.

To characterise the temporal structures of both the simulated data and neural data, we adapted the measures of bursty-ness and memory of Goh and Barabasi [3]. While the bursty-ness coefficient, *b* defined as

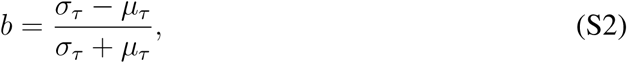

measures the broadness of the underlying ISI distribution with *μ_τ_* and *σ_τ_* as the mean and standard deviations of the interspike intervals respectively, the memory coefficient, *m* defined as

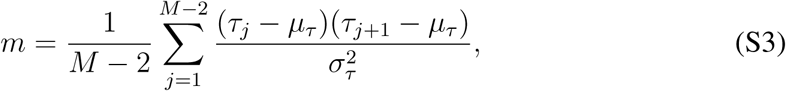

measures the short-time correlation between events.

For the stretched exponential distribution in Eq. (S1), the mean and standard deviations could be computed as

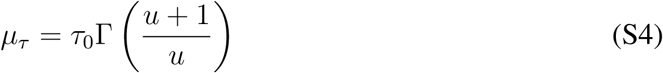

and

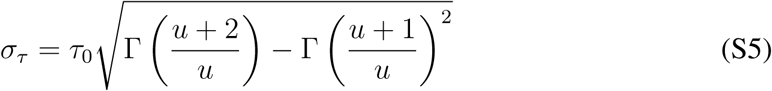

where Γ(*x*) ≡ (*x–*1)! is the gamma function. With these closed-form relationships, we could now study the limiting properties of the burstiness and memory coefficients. For Poisson processes, the mean, *μ_τ_*, and standard deviation, *σ_τ_*, coincide, i.e. *μ_τ_* = *σ_τ_* = *τ*_0_, and thus with Eq. (S2), give *b* = 0. For broad distributions, *u <* 1 in Eq. (S1), *σ_τ_ > μ_τ_* which gives *b >* 0 and tends to approach *b →* 1 in the limit *u →* 0. On the other hand, for narrow distributions, *u >* 1 in Eq. (S1), *σ_τ_ < μ_τ_* resulting to *b <* 0 and tends to *b → –* 1 in the limit *u→∞*. Hence, the bursty-ness parameter, *b*, is a bounded parameter, i.e., *b* ∈ [–1, 1].

For the synthetic datasets, note that fixing the parameter *u* automatically fixes the bursty-ness coefficient, *b*. However, because the synthetic interspike intervals are sampled independently, the memory coefficient, *m*, is approximately zero. Short-term memory can then be introduced by first sorting the interspike interval in decreasing (or increasing) order which results to *m* ≈ 1. Randomly shuffling the ordered interspike intervals of a subset of interspike intervals (100 events at a time in this case) results to a monotonic decrease of *m*. In the limit of infinite data, the memory coefficient is bounded by [–1, 1]. These bounds may no longer hold in the case of limited data. Despite this, a positive memory coefficient indicates that a short interspike (long) interval between events tends to be followed by another short (long) interval and a negative memory coefficient indicates that a short (long) interspike interval between events tends to be followed by a long (short) interval.

With this, we found that the MSR increased with bursty-ness and memory for the synthetically generated dataset as seen in Figure S1A and B. We also sought to characterise the relationship between the number of events, *M*, with the MSR which can be addressed by changing the characteristic time constant, *τ*_0_, in Eq. (S1) wherein decreasing *τ*_0_ leads to more events and thus, increased log *M*. We found that MSR decreased with log *M* as seen in Figure S1C. This result is indicative that MSR of randomly generated events can be explained by log *M*.

Following the results on synthetic data, we also analysed temporal characteristics in real neural dataset. In the case of neurons in the mEC data, we also found that MSR decreased with the logarithm of the number of observed spikes, log *M*, as shown in Figure S1D. To determine how much of the calculated MSRs can be explained by the number of observed spikes, *M*, we linearly regressed MSR with log *M* shown as the dashed line in Figure S1D. Residuals were then calculated as the deviation of the calculated MSR from the regression line and thus, captures the amount of MSR that cannot be explained by log *M* alone. We showed in Figure S1E and F that the MSR for real dataset still contained information going beyond log *M* as the residual MSRs (with respect to log *M*) still retain the dependence with spatial and head directional information as already observed in Fig 2.

